# Membrane insertion of‐ and membrane potential sensing by semiconductor voltage nanosensors: feasibility demonstration

**DOI:** 10.1101/044057

**Authors:** Kyoungwon Park, Yung Kuo, Volodymyr Shvadchak, Antonino Ingargiola, Xinghong Dai, Lawrence Hsiung, Wookyeom Kim, Z. Hong Zhou, Peng Zou, Alex J. Levine, Jack Li, Shimon Weiss

## Abstract

We develop membrane voltage nanosensors that are based on inorganic semiconductor nanoparticles. These voltage nanosensors are designed to self-insert into the cell membrane and optically record the membrane potential *via* the quantum confined Stark effect, with single-particle sensitivity. We present here the approach, design rules, and feasibility proves for this concept. With further improvements, semiconductor nanoparticles could potentially be used to study signals from many neurons in a large field-of-view over a long duration. Moreover, they could potentially report and resolve voltage signals on the nanoscale.

---

Recent advances in inorganic colloidal synthesis methods have afforded the construction of functional semiconductor (SC) nanoparticles (NPs) with ever-increasing control over size, shape, composition, and sophisticated heterostructures that exhibit unique photophysical, chemical and electronic properties^1,2,3,4^. This precise command of nanoscale materials synthesis has allowed for the exquisite engineering of excited state wavefunctions^5,6,7^, charge confinement, spatiotemporal control of charge-separated states^8^, and manipulation of Fermi levels and redox potentials. As a result, SC NPs have proved to be very useful in numerous applications in optoelectronics^9,10^, biological imaging^11^, sensing^12,13,14^, catalysis^15^, and energy harvesting^16^.

Integrating inorganic nanomaterials with naturally evolved or synthetically evolved biological machineries could yield highly sophisticated hybrid nanobiomaterials that outperform biological-only or inorganic-only materials. Such materials could be self-assembled by biomolecular recognition while maintaining the superior properties of inorganic materials^17,18^. Self-assembly of inorganic components by biomolecular recognition could align components in defined geometries, spatial orientations, and structures. In addition, careful design and control of the organic-inorganic interface could afford hybridization of electronic states, enhancement of radiationless energy transfer or electron transfer, or matching of Fermi levels with redox potentials.

Numerous functionalization and bioconjugation methods have been developed for the integration of inorganic-biological hybrid nanomaterials that are water soluble and biologically active^19,20^. Such hybrid nanomaterials have been used for *in vitro* biosensing, intra-cellular biological imaging^21^, single protein tracking in live cells^19^, and *in vivo* molecular imaging with favorable *in vivo* biodistribution and targeting properties (including renal clearance)^11,22,23^.

Much fewer attempts have been made to functionalize nanomaterials in a way that will allow their integration into the membrane. The ability to impart membrane protein-like properties to NPs could afford their targeting and insertion into the lipid bilayer and the construction of membrane-embedded hybrid nanomaterials with useful functions. For example, a few attempts have been made to target and insert (very small, < 3 nm) SC quantum dots (QDs) into the lipid bilayer. Al-Jamal *et al*. incorporated very small QDs in between the two lipid layers of the vesicle’s bilayer, proved by fluorescence microscopy^24^. Kloepfer *et al*. reported the transmission electron microscopy (TEM) micrographic evidence of QDs inserted into vesicles’ membranes^25^. Gopalakrishnan *et al*. successfully delivered lipophilic QDs (that were first loaded to vesicles’ membranes) into membranes of HEK293 cells *via* vesicle fusion^26^. Wi *et al*. investigated the maximum allowed QDs’ size both experimentally and theoretically that could still afford membrane insertion^27^. Recently, insertion of other types of nanomaterials into the membrane was demonstrated. Synthetic ion channels made from DNA nanostructures^28,29^ and ion channels made from carbon nanotubes^30^ were successfully inserted into lipid bilayers while maintaining functional ion transport across the membrane.

Following works on asymmetric type-II seeded nanorods (NRs) at low temperature on the single particle level^6,31^ and at room temperature on the ensemble level^32^, we recently demonstrated that these NRs exhibit a large quantum confined Stark effect (QCSE) at room temperature on the single particle level^33^. We also performed calculations that predict voltage sensitivity that is high enough for recording action potentials with good S/N ratio when such NRs are embedded in the plasma membrane (*In-preparation*). No attempts, to the best of our knowledge, have been made, however, to target and insert rod shaped nanoparticles into the lipid bilayer. In particular, membrane insertion of NRs with length larger than the membrane thickness (~4 nm) has not been demonstrated thus far. We present here an approach for inserting and positioning NRs as well as QDs in the membrane by imparting them with membrane protein-like properties and report on membrane voltage sensing experiments with these NPs.

## Results

We rationalized that NPs in the shape of cylindrical NRs could be functionalized with transmembrane *α*-helixes (by ligand exchange) and imparted with membrane-protein-like properties *i.e*. with hydrophobic side surface and hydrophilic tips (top and bottom) that extrude on both sides of the membrane (Supplementary Fig. SI-1 and Fig. SI-7.1). To do this, we modified our previously developed peptide coating technique for solubilizing QDs in biological fluids^11,34,35,36,37,38^ by using a novel, *α*-helical peptide sequence designed to insert the nanorods in a vertical orientation into membrane. The peptide sequence contains several cysteines along one face of the helix, hydrophobic amino acids on the opposite face, and hydrophilic amino acids at the N and C peptide termini: Myristoyl-CLTCALTCMECTLKCWYKRGCRGCG-COOH (Lifetein, New Jersey, U.S.A). Circular dichroism spectroscopy showed that the peptides adopted *α*-helical structures (in a membrane-mimicking octanol solution, Supplementary Fig. SI-2)Upon exchange of ligands with the helical peptides (Material and Methods), we hypothesized that the long axis of the peptide will preferentially bind along the long axis of the NR and that ~8-12 peptides will self-assemble on a single NR (Supplementary Fig. SI-1c). To test this hypothesis, we engineered a biotin at the peptide’s N terminus: Biotin-CLTCALTCMECTLKCWYKRGCRGCG-COOH. After ligand exchange, NeutrAvidin and biotinilated spherical Au NPs (GNPs) were added. TEM micrographs of the self-assembled complexes clearly show that GNPs are preferentially (> 85% of all cases) bound to the tips of the NRs rather than on their sides (Figs. 1a and 1b and Supplementary Fig. SI-3), confirming our hypothesis.

**Fig. 1:**
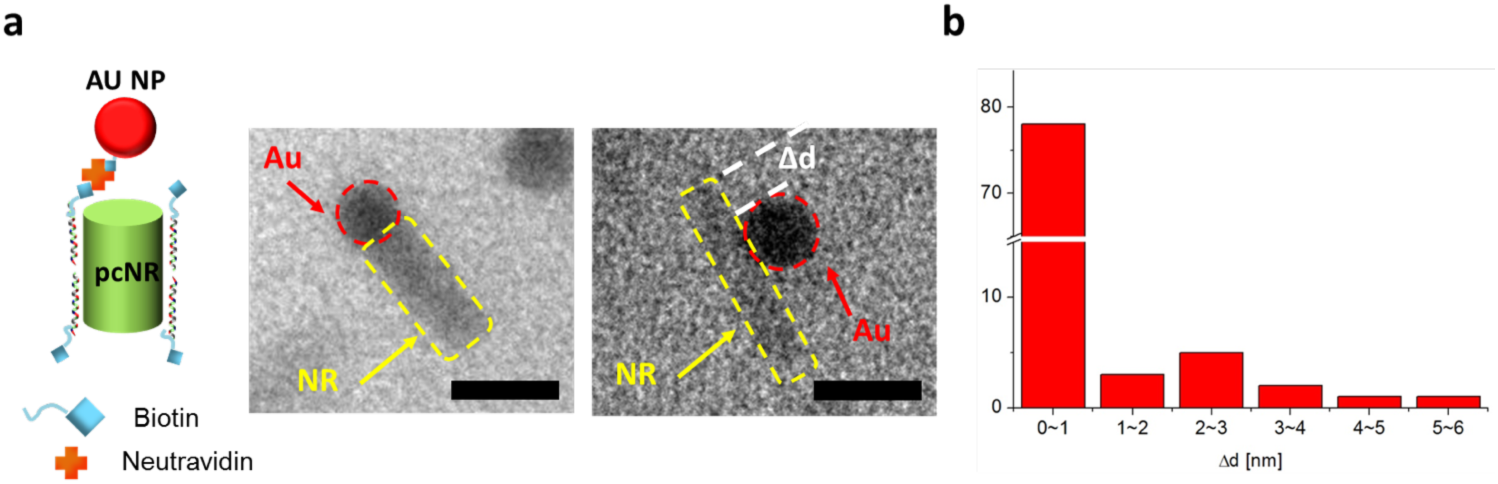
Characterization of peptides orientation in functionalized NRs. **(a)** TEM micrographs of biotinlated-GNPs bound to peptide-coated NRs (linked via NeutrAvidin) proving that the long axis of the peptide is parallel to the long axis of the NR. Scale bars are 10 nm. **(b)** Histogram of relative GNP position (Δd), measured from the tip of the NRs.

Although the peptide was designed for coating NRs, we also applied it to QDs, which also exhibited association with the membrane. To test the ability of peptide-coated NRs (pcNRs) and peptide-coated QDs (pcQDs) to self-insert into membranes, they were added to a preparation of electro-swelled giant unilamellar vesicles (GUVs). Rapid and spontaneous association of pcNRs with the membrane was observed, as confirmed by confocal microscopy (Fig. 2 and Supplementary Figs. SI-4.1 and 4.2). Moreover, when GUVs were composed of fusogenic lipids,pcNRs could be transferred to the plasma membrane of live cells by vesicle fusion (Figs. 2b, 2c and Supplementary SI-4.1). Here, we used the fusogenic lipids 1,2-stearoyl-3-trimethylammonium-propane (DOTAP) or 3β-[N-(N’,N’-dimethylaminoethane)-carbamoyl] cholesterol hydrochloride (DC-Chol) mixed with a cone shaped lipid, 1,2-dioieoyi-.*sn*-giycero-3-phosphoethanolamine (DOPE)^26,39,40^. Fig. 2b captures the moment of fusion of pcNRs loaded fusogenic vesicles with a HEK293 cell (bright field) while Fig. 2c shows a subsequent fluorescence image after the fusion event. A z-stack of the same cell is shown in Supplementary Fig. SI-4.1, clearly indicating membrane staining.

**Fig. 2:**
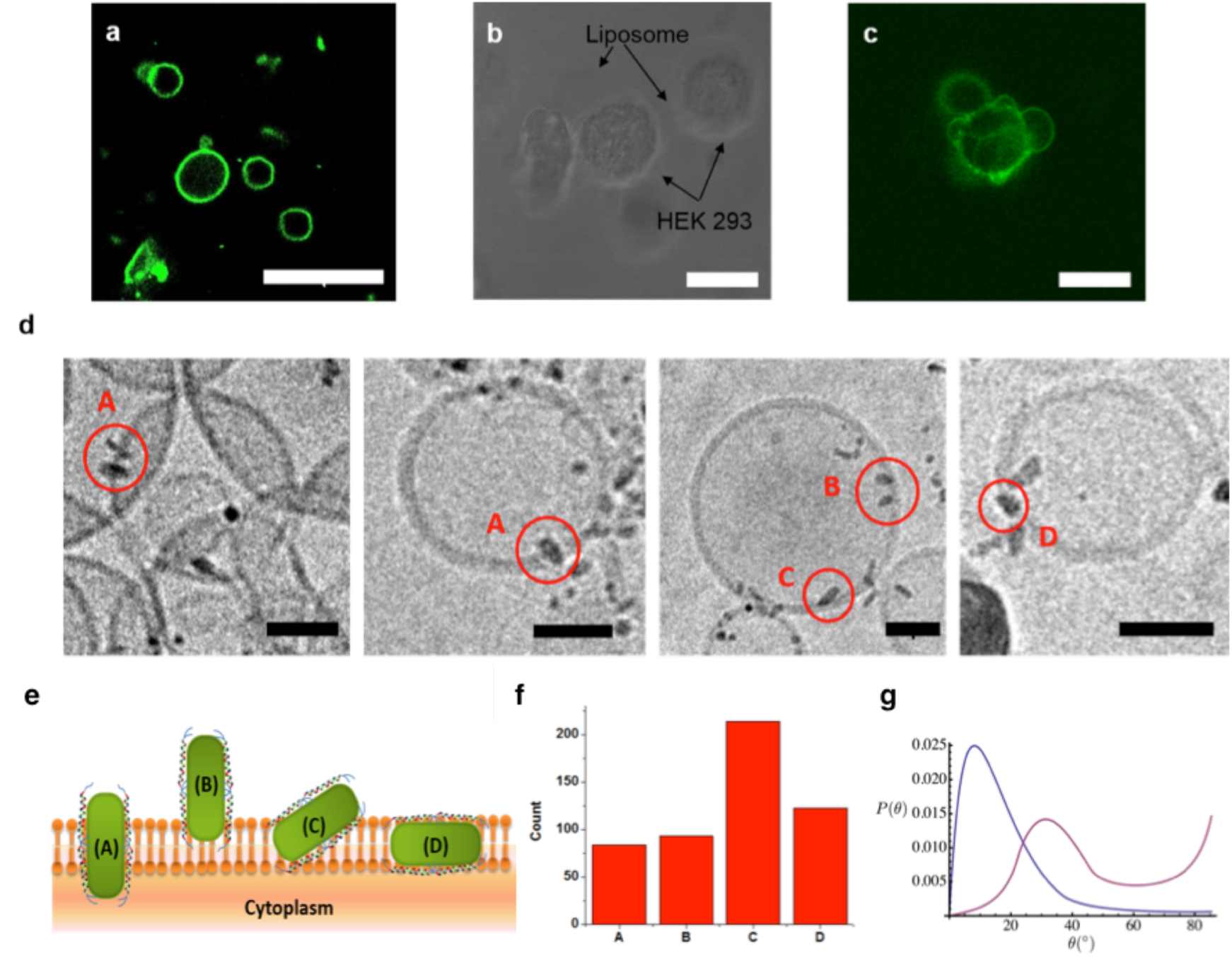
Demonstration of membrane insertion of functionalized NRs. **(a)** NRs loaded GUVs **(b,c)** Vesicle fusion to HEK293 cell (**b:** bright field, **a, c:** confocal) **(d)** CryoEM micrographs of pcNRs inserted into SUVs. **(e)** Schematics of possible pcNRs insertion geometries or attachments to the lipid bilayer: (A) properly inserted (B) partially inserted (C) attached in an angle (D) horizontally embedded. **(f)** Histogram of insertion geometries (A) – (D). (Scale bars are 10 *μ*m (a~c), 30 nm (d)) **(g)** Model calculations (see Supplementary SI-7) of canting angles (θ) probability distribution for a membrane-inserted nanorod. Calculations for no hydrophobic mismatch (L = t = 4 nm, blue) and for significant hydrophobic mismatch (L = 6 nm, t = 4 nm, red) are shown. In both cases the rods are terminated at both ends by hydrophilic cylinders of length 2 nm (details of the model are discussed in Supplementary SI-7).

The orientation of the membrane-associated pcNRs was probed by polarization microscopy^41^, capitalizing on the fact that NRs’ absorption and emission dipoles are aligned along their long axis^42^. To estimate orientation of pcNRs in the membrane, we imaged the fluorescence of GUVs loaded with pcNRs using linearly polarized excitation. Since the absorption and emission dipoles of NRs are aligned along their long axis, polarized excitation could determine if pcNRs are aligned parallel or perpendicular to the membrane’s lipids (Supplementary Fig.SI-5). By analyzing the polarization anisotropy of individual pcNRs and fitting to a simple model), we could estimate that ~ 58% of the pcNRs were inserted with an orientation that is parallel to the lipid molecules (perpendicular to the membrane plane, Supplementary SI-5), supporting the notion of membrane insertion.

To further investigate the level of association and/or membrane insertion, small unilamellar vesicles (SUVs) loaded with pcNRs were flash-frozen and imaged by cryoelectron microscopy (cryoEM) (Fig. 2d). SUVs were first sonicated and then extruded through a 100 nm pore filter. Since cryoEM images are 2D projections, exact z-positions of pcNRs are not exactly known. For this reasons, the level of insertion of pcNRs was assessed only for particles close to the vesicle’s ‘equator’. We analyzed over 500 pcNRs and classified them into the four categories (Fig. 2e and 2f). A-type represents an ideal, symmetric and perpendicular insertion (the NR symmetrically traverses the membrane), it was observed for 16.4% of the pcNRs. B-type represents partial (asymmetric) but perpendicular insertion (18%). C-type represents partial, tilted insertion and is the most abundant (41.7%of the pcNRs). D-type represents horizontal insertion in between the two leaflets of the membrane (23.9%). The histogram in Fig. 2f shows the partitioning in insertion geometries. Because of the projection and ambiguity in z positioning, these percentages are not precise estimate for the partitioning between the different insertion configurations. Nonetheless, despite the fact that we have tested thus far only one rationally designed *α*-helical peptide sequence, a sizeable fraction of pcNRs showed proper membrane insertion, proving the feasibility of this functionalization approach. A control cryoEM experiment showed that as-synthesized (ligand-coated) NRs do not insert into vesicles’ membranes (Supplementary SI-6).

To assess the cryoEM results, we developed a statistical mechanics model that calculates the pcNR membrane insertion energy and insertion (or canting) angle (θ). This model predicts that with hydrophobic surfaces covering a length of the pcNRs comparable to the membrane thickness, the fraction of rods inserted into the membrane approaches unity in thermal equilibrium. In order to stabilize the orientation of rods in the membrane to be close to the membrane’s normal, it is advantageous to include hydrophilic ends on the nanorods. For reasonable lengths of these ends,they do not significantly change the partitioning of rods between the membrane and the solvent. Moreover, the model predicts a canting angle distribution (Fig. 2g) that resembles the histogram in Fig. 2f, suggesting some degree of hydrophobic mismatch (Supplementary Figs. SI-7.1 and SI-7.2). We note, however, that the statistical nature of ligand exchange with the designed peptide does not necessarily impart precise hydrophobic surfaces and hydrophilic tips.

Delivery and insertion of pcNRs to the plasma membrane of live cells are not restricted to GUV or SUV fusion. pcNRs could also be added directly to the growth media of a tissue culture, as demonstrated in Fig. 3 for HEK293 cells. By diluting the concentration of pcNRs, sparse labeling could be achieved such that individual (or small aggregates of) pcNRs could be clearly observed (Fig. 3c, red arrow and Supplementary SI-8.1). The intensity trajectory of the pcNR marked with a red arrow in Fig. 3c (Fig. 3d) exhibits clear blinking (intermittency) in its emission pattern, which is indicative of bright emission from a single particle^43^.

**Fig. 3:**
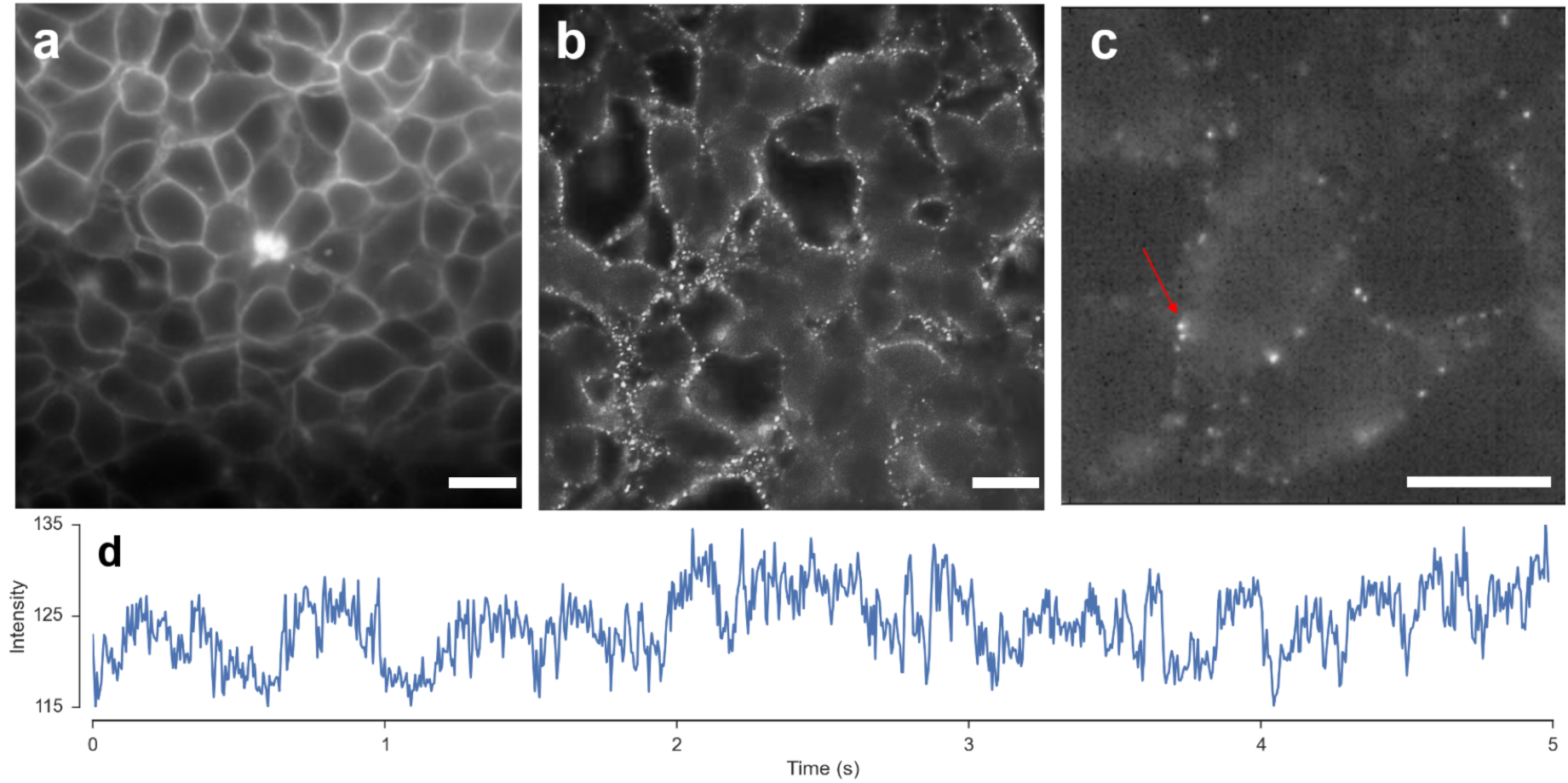
Demonstration of membrane insertion of functionalized NRs into HEK293 cells. Fluorescence images of HEK293 cells stained with **(a)** di-8-ANEPPS (control) and **(b, c)** pcNRs targeted to membranes at high **(b)** and low **(c)** concentrations. Red arrow in **(c)** points to a single blinking pcNR whose intensity time-trace is shown in panel **(d)**. Scale bars are 10 μ,m.

Not only these pcNRs are bright, but theoretical investigations also predict that when embedded in the membrane, they could exhibit superior voltage sensitivity as compared to conventional voltage sensitive dyes^44^. To test these predictions, we attempted to measure the voltage response of membrane inserted peptide coated nanoparticles using two different approaches.

In the first, pcQDs were inserted into membranes of engineered self-spiking HEK293 cells^45^ and fluorescence signals were recorded. Fig. 4 shows optical recordings of self-spiking HEK293 cells with pcQDs. By averaging the fluorescence signals from all pixels in a movie frame (for all frames) and applying a simple threshold, a digital map of membrane pixels stained with pcNPs’ is obtained (Fig. 4b). Fig. 4c shows a Δ*F/F* time-trace (averaged over all white pixels in Fig. 4b as function of time), clearly capturing coherent voltage oscillation (only the modulated part of the emission is shown). Voltage oscillations are also detected from single points (average over 5 x 5 pixels) on the membrane as shown for 3 different points per cell for 3 different cells (Figs. 4e-4g). These single point trajectories are noisy due to unstable, dynamic fluctuations in membrane insertion, uncorrelated blinking^43^, and different/fluctuating dielectric and electric environments. A cross-correlation (Fig. 4d) between recordings from red and green pixels (in Fig. 4a) shows a significant correlation between time points that are < 6 s apart, which attests to the oscillatory nature of collective self-spiking signal and reports on the de-coherence time (and length) between cells. Voltage oscillations are found for all recorded points. Fig. 4h shows an overlay of the 9 trajectories in Figs. 4e-4g where the asterisks in Fig. 4h point to differences in oscillations phase (phase of green points precedes the phase of red points). In a separate experiment, to assess if the time traces shown in Fig. 4 indeed report membrane voltage oscillations, tetrodotoxin (TTX), a toxin that blocks the sodium ion channel (and therefore inhibits action potentials in neurons^45^) was added to the cell culture during the optical recording. As in the control experiment (using a commercial voltage sensing dye ANEPPS, blue trajectory in Fig. 4i), oscillations in trajectories recorded from cells labeled with pcQDs diminish within ~5 seconds after the addition of TTX (black trajectory in Fig. 4i).

**Fig. 4:**
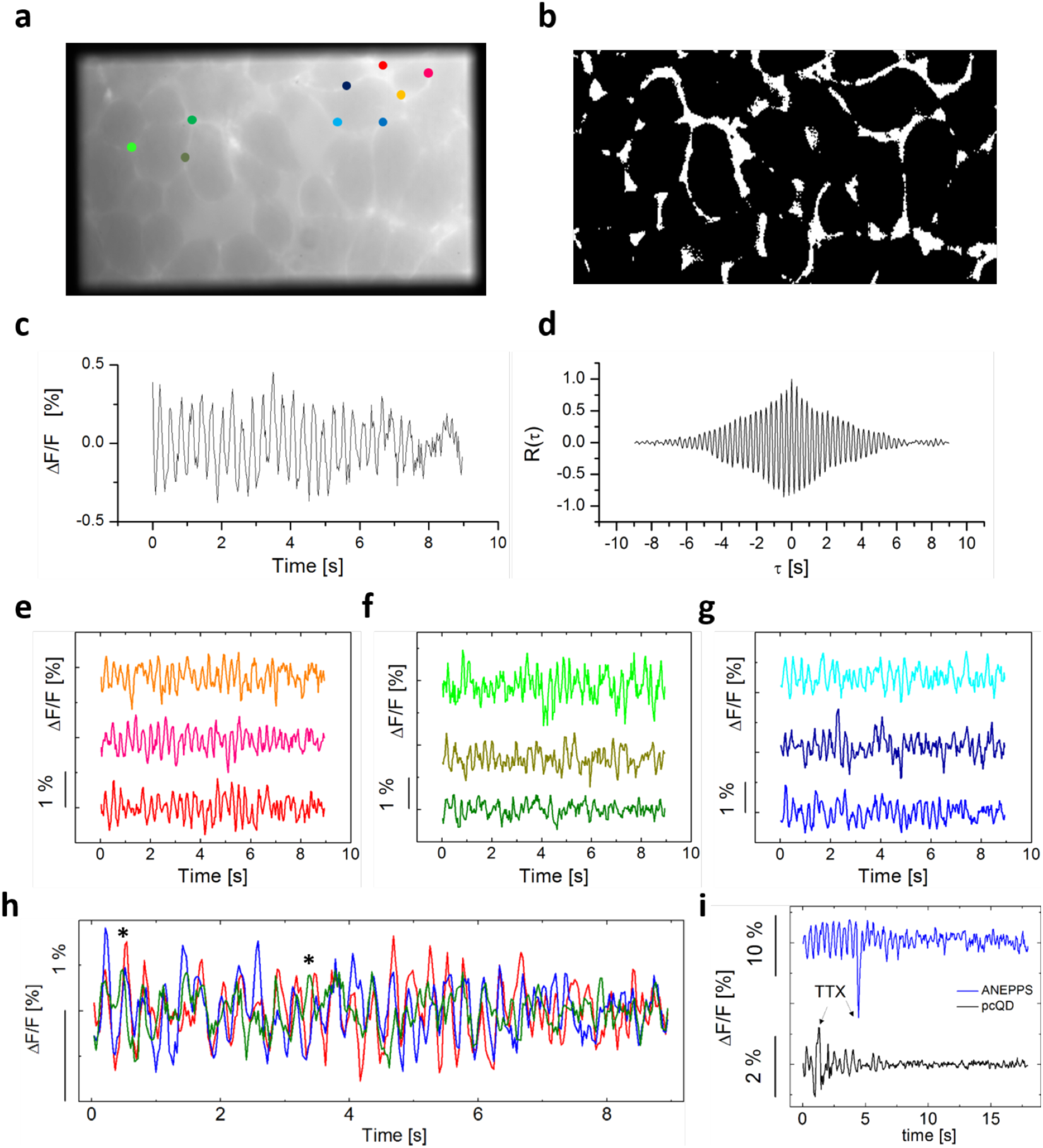
Simultaneous multisite voltage sensing in self-spiking HEK293 cells with pcQD. **(a)** Imaged field-of-view. **(b)** Segmentation of membrane pixels in (a). **(c)** Δ*F/F* time-trace (averaged over all white pixels in (b); DC component not shown). **(d)** Temporal cross correlation of red and green cells signals in (a). **(e~g)** Non-averaged Δ*F/F* time-traces at different points as indicated in (a). **(h)** Comparison of non-averaged Δ*F/F* time-traces between three points at red (e), green (f), and blue (g). **(i)** Non-averaged *AF/F* time-traces for ANEPPS (control) and pcQDs after addition of TTX toxin (that blocks the Nav1.3 sodium ion channel).

In the second approach, pcNRs were inserted into membranes of wild-type HEK293 cells with membrane potential modulated *via* whole-cell patch clamp. Fluorescence emission and membrane voltage were recorded simultaneously. For these experiments, non-ideal quasi type-II CdSe-CdS pcNRs (sample #3 in ref. 33) of dimensions 4 × 10 nm (diameter × length) were used (the synthesis of more suitable true type-II NRs with higher voltage sensitivity and proper dimensions is currently being pursued). pcNRs were applied directly to wild-type HEK293 cells cultured on a coverslip. Fluorescence movies were recorded in synchrony with the membrane voltage modulation (with a cycle of 2 movie frames recorded at ‐150 mV followed by 2 movie frames recorded at 0 mV, voltage modulation frequency of 100 Hz and duration of 2000 frames).

Fig. 5a shows a fluorescence time trajectory recorded from a single (or possibly a small aggregate of) pcNR(s) (as judged by blinking; highlighted by an arrow in Fig. 3(c). A link to the movie is provided in Supplementary SI-8.1). The fluorescence trajectory is highly noisy, most likely due to fluorescence intermittency (blinking) and unstable, dynamic fluctuations in membrane insertion (see discussion about membrane insertion stability in Supplementary SI-7). A zoom-in to the trajectory at around 4.6 sec (Fig. 5b) shows a zig-zag pattern in the fluorescence intensity that is synchronized with the modulated clamped voltage. We defined, for each semi period, the signal Δ*F/F* as the difference between voltage-on and voltage-off intensities divided by the mean time-trace intensity. This signal (Δ*F/F*) exhibits a high degree of variations throughout the acquisition time (5 s) with a few spikes of high signal about 100 ms long.

We identified 8 individual (or small cluster of) pcNRs in the patched cell’s membrane (or its proximity) and computed the mean signal during the entire time-trace (excluding the off periods of the pcNR fluorescence blinking) for each. Out of 8 pcNRs, only 3 exhibited a mean absolute signal that was higher than the mean calculated for pcNRs in membranes of non-patched cells.These 3 pcNRs had a negative mean signal (with a ±1σ error range that does not include 0), while the control group of pcNRs in the membrane of non-patched cells exhibited a mean signal which is statistically indistinguishable from 0 (Supplementary Fig. SI-8.2 and SI-8.3).

Since the signal Δ*F/F* exhibits spikes or “bursts” of high signal that presumably correspond to brief periods of membrane insertion, we carried out an objective analysis that is focused on such brief “bursts”. For this purpose, we performed a running average of the Δ*F/F* signal and identified bursts for which the average is higher than a threshold. We then computed the sum signal in each burst *i* as: 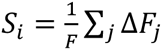, which we dub as the ‘burst score’ (see Materials and methods and Supplementary SI-8 Patch-clamp analysis). Fig. 5c shows the distribution of burst scores for bursts belonging to pcNRs found in the patched cell membrane (patched in-phase, red) compared to burst scores of controls that should not exhibit any correlated signal. In particular, in the out-of-phase controls, we suppress any intensity fluctuation in-phase with the voltage modulation by averaging frames corresponding to ON and OFF voltage semi-periods (see Materials and methods). We observe that while the controls exhibit an even distribution of bursts with positive and negative scores (consistent with random fluctuations), bursts of patched pcNPs show a predominance of negative scores (consistent with fluorescence reduction induced by the applied voltage). Note, however, that only 18 in-phase bursts were identified and analyzed for the 3 pcNRs associated with the patched cell membrane (and 20 out-of-phase bursts). For the non-patched cells control we identified 7 pcNRs (in non-nearest neighbor cells) yielding a total of 28 in-phase bursts (and 40 out-of-phase bursts).

**Fig. 5:**
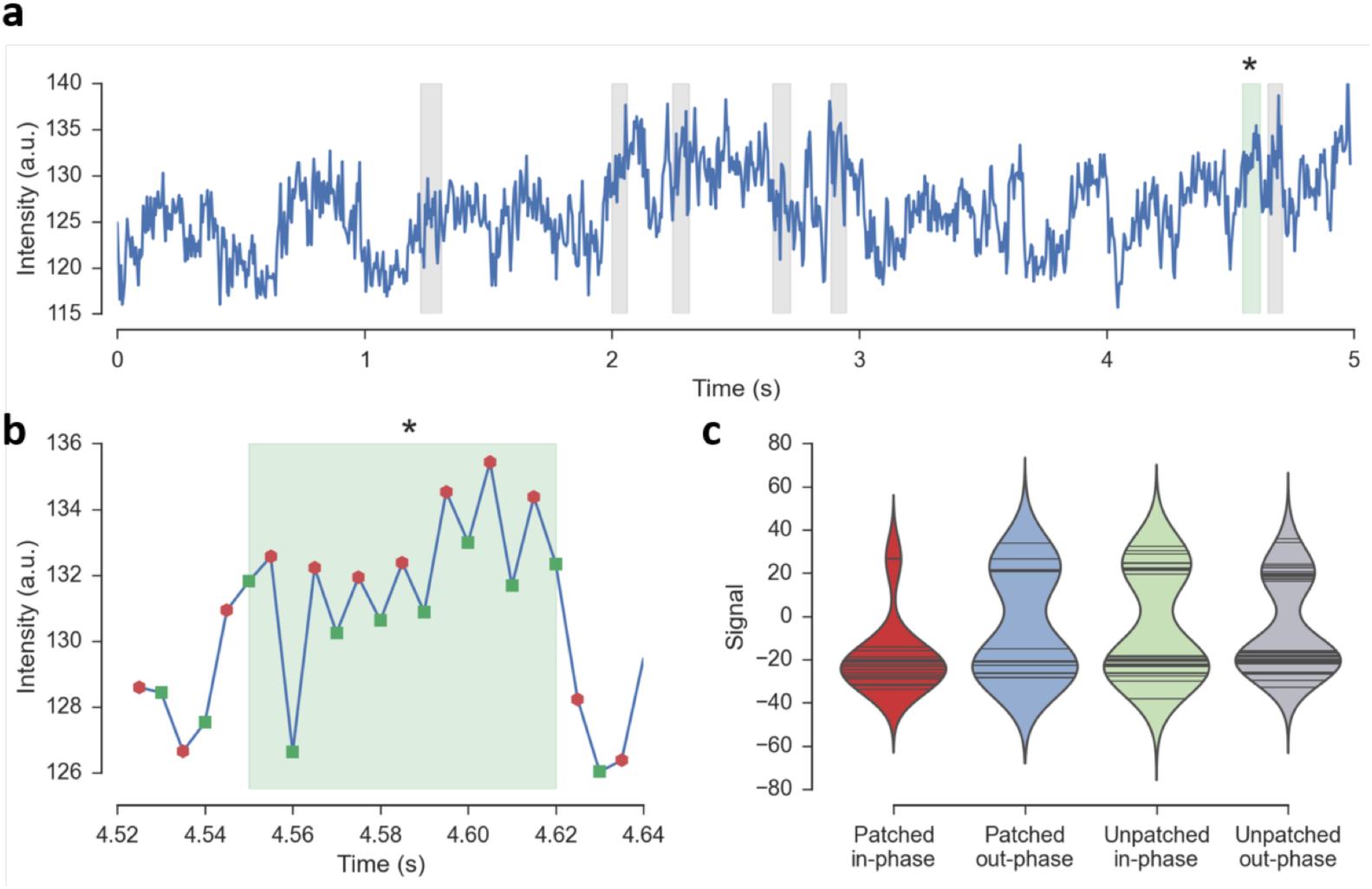
Voltage response of pcNRs. **(a)** Intensity trace with highlighted (shaded) intervals of large ΔF/F signals (“bursts”). **(b)** Zoom-in intensity trace of a burst in (a); each marker represents a 2-frame average intensity during the voltage-on (*green squares*) and voltage-off (*red dots*) semi-period. **(c)** Distribution of burst scores for all the bursts in a video. The first group (*red*) represents patched pcNRs that exhibit the highest signal. The other 3 distributions are control groups for unpatched pcNRs and/or out-of-phase signals.

Overall, our results suggest that membrane inserted pcNRs exhibit membrane voltage sensitivity, but this conclusion is stated with great caution due to the high noise level and the limited number of pcNRs in the membrane of the patched cell.

The data presented in figures 4 and 5 suggest that with further improvements, pcNRs could be possibly used in the future for membrane potential recording. The signal quality could be greatly increased by a series of enhancements. For example, preliminary experiments and calculations suggest that seeded nanorods heterostructures with type-II band offset and large seed position asymmetry could exhibit very high voltage sensitivity^33^. Moreover, improved membrane insertion stability will reduce measurement noise and enhance the signal. Lastly, as previously shown^33^, shifts in spectral peak position are considerably more sensitive than Δ*F/F* changes. Simple modification to the optical set-up (based on “dual-view” microscopy^46^) could enhance voltage sensitivity even further.

Development of high sensitivity pcNRs could afford unprecedented ways for studying electrical activities in neuronal, neuromuscular, and visual systems, offering superresolution voltage sensing on the nanoscale (such as across a single synapse), or the ability to record large number of signals from a large-field of view (high throughput recording). pcNRs could also find applications in other areas of science and engineering, as for example, in inducing action potential^47,48^, characterization of high-density fast integrated circuits and energy harvesting by membrane-inserted artificial light harvesting complexes. Lastly, the ability to impart membrane protein-like properties to inorganic and organic nanoparticles could allow the construction of novel membrane-based hybrid (organic-inorganic) materials with unique exploitable properties.

## Acknowledgements

We acknowledge the generous help of Dr. Adam Cohen, for providing the self-spiking HEK cell line, and for providing access to his laboratory and patch-clamp fluorescence set-up. We also acknowledge the use of instruments at the Electron Imaging Center for NanoMachines supported by NIH (1S10RR23057 to ZHZ) and the Advances light Microscopy and Spectroscopy core, both at the Caifornia Nano Systems Institute (CNSI) at UCLA. SW acknowledges funding from BSF #2010382 and DARPA #141-002 #D2133. This material is based upon work supported by the U.S. Department of Energy Office of Science, Office of Biological and Environmental Research program under Award Number DE-FC02-02ER63421. AJL acknowledges partial support from NSF-DMR-1309188.

## Author contribution

KP performed experiments, data analysis, and helped writing the paper

YK performed experiments, data analysis, and helped writing the paper

VS designed the peptides and helped editing the paper

AI performed signal analysis for the patch clamp experiments

XD performed the vesicle imaging with cryoEM.

LH performed experiments.

WK performed experiments.

HZ performed the vesicle imaging with cryoEM.

PZ performed the patch-clamp experiment.

JL synthesized NPs for this study

AJL developed the theory and performed simulations for membrane insertion.

SW designed and managed the project and helped writing the paper

## Material & Methods

### NR Synthesis

Quasi-Type-II NRs (CdSe seeded in CdS):Cadmium oxide (CdO, 99.99%), tri-n-octylphosphine (TOP,90%), trioctylphosphine oxide (TOPO, 99%), selenium (Se, 99.999%) and Sulfur (S, 99.5%) along with all organic solvents were purchased from Sigma-Aldrich and used without any further purification. Hexylphosphonic acid (HPA) and octadecylphosphonic (ODPA) were purchased from PCI Synthesis.

A 50 ml round bottom flask was loaded with 60 mg (0.5 mmol) CdO, 280 mg ODPA and 3 g TOPO. After degassing under vacuum for 1 hr at 120 °C the temperature was raised to 340 °C under argon until dissolution of CdO at which point 1.8 ml TOP was injected and temperature was raised to 370 °C. A solution containing 58 mg Se in 0.5 ml TOP was swiftly injected and heating mantle was removed. Final core size had a diameter of about 2.7 nm. A slight modification of previously reported methods^7^ was used for seeded growth of CdS. A 50 ml round bottom flask was charged with 211 mg (1.6 mmol) CdO, 1 g ODPA, 50 mg HPA and 3.46 g TOPO. The reaction flask was degassed for 3 hrs at 130 °C and then temperature was raised to 340 °C under argon until dissolution of CdO at which point 1.8 ml TOP was injected. CdSe seed solution was separated and purified for reaction by mixing with toluene and precipitating with excess methanol 3 times. Seeds were then re-dissolving in 0.6 ml TOP. The S:TOP precursor solution was prepared by mixing 51 mg S (1.6 mmol) in 0.6 ml TOP. Temperature was raised to 350 °C for injection. The amount of dots used was 8×10^-7^ moles.

### NR functionalization with peptides

The sequences of the two peptides used in this study are:

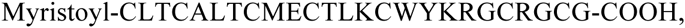

and:

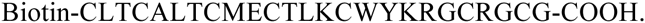

Peptides were purchased from LifeTein LLC, purified to a level of 70 % by HPLC, characterized by mass spectrometry and circular dichroism (Supplementary Fig. SI-2). The protocol for NRs functionalization with *α*-helical peptides is similar to the protocol reported in^35^ with the following modifications: as-synthesized NRs were coated with hydrophobic surfactants such as TOPO or ODPA. To exchange these surfactants with the designed peptides, we first stripped the surfactants off the NRs by multiple (5~6×) methanol precipitation steps, followed by redissolution in pyridine 450 μl. The NR’s concentration was 0.1 μM. 4.0 mg of peptides were dissolved in 50 μl of DMSO, and mixed with NRs in DMSO solution. 12 μl of tetramethylammonium hydroxide (TMAOH) were added to the solution to increase the pH to 10.0 allowing the peptides to bind to the NRs’ surface efficiently. The mixture was then centrifuged and redispersed in 150 μl of DMSO, in a form ready to be used for cell membrane insertion (staining). For vesicle staining or for cryoEM experiment, NRs in DMSO solution were eluted through a G-25 Sephadex desalting column (Amersham, Piscataway, NJ) and equilibrated with PBS buffer. The pcNRs were stored at 4 °C.

### GNP-pcNR complex

Biotinylated gold (Au) nanocrystals were a gift from Ocean NanoTech Inc. CdSe/CdS NRs were functionalized with biotinilated peptides using the same protocol as described above. 1 μl of the biotinilated peptide-coated NRs solution was mixed with 15 μl of neutravidin solution (2 mg/ml) and incubated for 20 mins at room temperature. 1 μl of 20X diluted biotin-Au solution was then added to this mixture and incubated for 2 hrs. 2 μl of the final solution was then deposited on copper EM grids and let dry in air.

### Loading pcNRs into vesicles

1,2-dimyristoyl-sn-glycero-3-phosphocholine(DMPC),1,2-dioleoyl-3-trimethylammonium propane chloride salt (DOTAP), 3β-[N-(N’,N’-dimethylaminoethane)-carbamoyl]cholesterol hydrochloride (DC-Cholesterol), were purchased from Avanti Polar Lipids, Inc. Chloroform solutions of DOTAP (25mM, 6μl), DMPC(10mM, 6μl) and DC-Cholesterol (10mM, 6 μl) were mixed and dried in a vacuum for 4 hrs in a rotary evaporator. The film was then hydrated with 1 ml of 0.1M sucrose containing PBS buffer with pH 6.24 overnight at 37°C incubator, during which vesicles were spontaneously formed. Vesicles were stored at 4 °C unless used in experiments (they are stable and useable for about one week). For fluorescence microscopy measurement, 2 μl of pcNRs (eluted through a G-25 Sephadex desalting column) were added to the 10 μl of vesicle solution. They spontaneously and rapidly (~1 min) self-inserted into the vesicles’ membranes.

For GUVs, the same lipid composition (6 μl of 25 mM DOTAP, 6 μl of 10 mM DMPC, and 6 μl of 10 mM DC-Cholesterol) was diluted with 200 μl of chloroform. 50 μl of the lipid in chloroform solution were loaded on the indium-tin-oxide (ITO) coated glass. After 30 minutes of dry, the other ITO coated glass was faced to the lipid dried ITO glass. Two glasses are separated by the 3 mm thickness of O-ring, forming the aqueous chamber for electroswelling. 10 Hz of 1.0 V square voltage pulse was applied to the two ITO glasses for 20 mins, followed by GUV preparation for imaging.

### Cryoelectron microscopy

For cryoelectron microscopy measurement, 10 μl of pcNRs (eluted through a G-25 Sephadex desalting column) were added to 50 μl of vesicle solution. An aliquot (3 μl) of sample was placed on holey carbon-coated Quantifoil grid, manually blotted with filter paper, and plunged into liquid ethane to make a cryoEM grid with vesicles embedded in vitreous ice. The grid was transferred to a Gatan 626 cryo sample holder cooled down by liquid nitrogen, and inserted into an FEI TF20 cryoelectron microscope for imaging at 200 kV operating voltage. Images were recorded at several magnifications on a 4k x 4k CCD camera (TVIPS) at ~5 μm under focus with an accumulated electron dosage of ~20 e^-^/Å^2^ on each sample area.

### Cell culture and staining

HEK293 cells (AATC, VA) were maintained in 1:1 Dulbecco’s Modified Eagle Medium and Nutrient Mixture F-12 (Invitrogen, NY) supplemented with 10% fetal bovine serum (FBS; Sigma-Aldrich, MO), 0.6 mg/ml of Geneticin (G418) (Life technologies), and 5 μg/ml of puromycin (Life technologies). Cells were grown on 35 mm glass bottom dishes until they reached 90 % confluency. The same protocol was applied to self-spiking HEK293 cells. For ANEPPS staining, di-8-ANEPPS solution in DMSO was added directly to the cells in a 35 mm glass bottom dish to a final concentration of 0.1 μM. Cells were then incubated at 4 °C for 5 mins before imaging.

### Optical imaging and data acquisition of pcQDs’ fluorescence signal in self-spiking HEK293 cells

The microscope set-up is based on an Olympus IX71 inverted microscope equipped with a Xenon lamp (Olympus, U-LH75XEAPO, 75W) and excitation filter (BP 470/40, Chorma Technology Corp, Bellows Falls, VT). The emission of the NRs was collected by a 60× objective lens (Olympus, Plan Apo 60×,n=1.45, oil immersion), passed through a dichroic mirror (DM, 505DCXRU, Chorma Technology Corp, Bellows Falls, VT). Imaging was done with an Andor iXon electron multiply (EM) charge coupled device (CCD) camera (EMCCD, Andor iXon, South Windsor, CT). 2 μl pcQDs in DMSO solution (~300 nM) were loaded to the glass-bottom dish (Fisher Scientific) where the self-spiking HEK293 cells were cultured in. The pcQDs spontaneously inserted into cell membranes within 1-2 mins. The pcQD loading density estimated from the image is ~10^5^ pcQDs per cell. After rapid shaking, the cell medium was changed with Dulbecco’s Phosphate-Buffer Saline (DPBS, Life technologies). The dish was then placed on the microscope. Fluorescence was recorded in a movie format for 9 seconds with a 30 ms integration per frame.

### Simultaneous patch-clamp recording and fluorescence imaging

2 μl of pcNRs were added directly to the cell culture (in a 35 mm glass bottom dish with 2 ml of cell culture medium). Cells were then incubated at 37 °C for 5 mins before patch-clamp and imaging. As estimated from images, about ~ 10 particles were inserted into each cell on average. The loading density is approximated to be ~ 10^-7^ pcNRs/ nm^2^.

All imaging and electrophysiology were performed in Tyrode’s buffer (containing 125 mM NaCl, 2.5 mM KCl, 3 mM CaCl2, 1 mM MgCl2, 10 mM HEPES, 30 mM glucose, at pH 7.3, and adjusted to 305-310 mOsm with sucrose). For patch clamp, filamented glass micropipettes (WPI) were pulled to a tip resistance of 5–10 MΩ, and filled with internal solution containing 125 mM potassium gluconate, 8 mM NaCl, 0.6 mM MgCl2, 0.1 mM CaCl2, 1 mM EGTA, 10 mM HEPES, 4 mM Mg-ATP, 0.4 mM Na-GTP (pH 7.3); adjusted to 295 mOsm with sucrose. Pipettes were positioned with a Sutter MP285 manipulator. Wholecell, voltage and current clamp recordings were acquired using a patch clamp amplifier (A-M Systems, Model 2400), filtered at 5 kHz with the internal filter and digitized with a National Instruments PCIE-6323 acquisition board at 10 kHz.

Simultaneous whole-cell patch clamp recordings and fluorescence recordings were acquired on a home-built, inverted epifluorescence microscope equipped with a 60× water immersion objective, numerical aperture 1.20 (Olympus UIS2 UPlanSApo 60×/1.20 W) and a scientific CMOS camera (Hamamatsu ORCA-Flash 4.0). 488 nm laser (Coherent Obis 488-50) intensity was modulated with an acousto-optic tunable filter (AOTF; Gooch and Housego 48058-2.5-.55-5W). Imaging of pcQDs was performed at illumination intensities of ~ 1 W cm^-2^. A long-pass dichroic filter (Chroma zt505-515+650NIR Tpc) and a band-pass emission filter (525/30) were used for fluorescence imaging. For fast data acquisition, a small field of view around the cell of interest was chosen at the center of the camera to achieve a frame rate of 1,000 frames per second.

### Data analysis of pcNRs’ fluorescence during patch-clamp recording

From the video we manually identify pcNP position on both the patched cell membrane and on non-patched cells. For each identified pcNP, the time trace of emission intensity {*t*_*k*_} is obtained by averaging for each frame *k* a circular region of roughly 20 pixels around the pcNP. The time trace intensity is binned each 2 frames in order to obtain an intensity {*t̅*_j_)} for each voltage alternation semi-period, then the difference {Δ*F*_i_} are computed as {(*t̅*_1_ — *t̅*_0_),—(*t̅*_2_ — *t̅*_1_),(*t̅*_*3*_ — *t̅*_*2*_),—(*t̅*_*4*_ — *t̅*_*3*_), …}. (the sign alternates and is “+” for ON-OFF and “-” for OFF-ON transitions). Finally these differences are divided by the average time trace intensity to obtain the signal {Δ*F*_*i*_/*F*}.

The burst search is performed as follows. The square of the running average of the {Δ*F*_*i*_/*F*} signal is computed and the time periods where this squared average is higher than a threshold (set to 60% of the maximum) are identified as bursts. Next, for each burst *i*, we extracted the total signal(*burst score*) 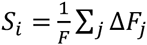. The out-of-phase case is obtained by removing the first video frame and applying the same analysis on the time traces. In this case, the binning step averages frames between ON and OFF semi-periods, suppressing any signal in-phase with the voltage alternation. See Supplementary SI-8 for detailed description of the patch-clamp data analysis.

